# Decoding of frequency-modulated sweeps by core and belt neurons in the alert macaque auditory cortex

**DOI:** 10.1101/2025.04.30.651560

**Authors:** Brian J. Malone, Gregg H. Recanzone

## Abstract

Acoustic stimuli where the spectrum is time-varying are ubiquitous in natural sounds, including animal vocalizations, human speech, and music. Early studies of such stimuli involving frequency-modulated sweeps revealed that neurons in the primary auditory cortex of a variety of mammals show differences in firing rates depending on either the direction of the sweep and/or the sweep velocity. Psychophysical studies have also shown that the perception of such time-varying stimulus parameters is quite acute, underscoring the importance of such signals in normal acoustic perception. Surprisingly, the responses of auditory neurons in alert primates has been little studied, and we have limited information relating neural activity to the perception of these signals. In this study, we investigated the neural discriminability of sweep direction and velocity for frequency-modulated sweeps presented to alert rhesus macaque monkeys in both core and belt auditory cortical areas. We quantified how well these information-bearing parameters were encoded using spike train pattern discriminators, and compared decoder performance when neural responses were restricted to temporal patterns or firing rates. Decoding accuracy for firing rate alone exceeded chance, and rate-normalized, spike-timing information was essentially equivalent to the complete firing pattern. Although most belt areas showed small decreases in decoding accuracy relative to the primary field, all fields encoded and represented sweeps similarly. Thus, there was little evidence of hierarchical processing between core and belt fields for these stimuli, indicating that frequency modulation sweep direction and velocity are not specifically extracted in the early auditory cortical hierarchy.

**Significance Statement:** Frequency modulated (FM) stimuli are a key feature of many time-varying acoustic stimuli, including speech, vocalizations, music, and environmental sounds. The direction and velocity of FM stimuli are major information-bearing parameters that allow one to discriminate and perceive these sounds. We tested whether single neurons in core and belt auditory cortical fields in alert macaque monkeys preferentially process these features along the cortical hierarchy. We found that the timing of neural activity was much more important than the absolute amount of activity in all cortical areas, and did not observe any evidence of improved discriminability in core or belt fields beyond that seen in the primary auditory cortex (A1).

## Introduction

Changes in the sound frequency over the course of a sound—frequency modulation—are ubiquitous in animal communication, human speech, music and other natural sounds. In laboratory experiments, frequency-modulated (FM) sweeps are often used to parameterize this aspect of natural sounds, and those experiments suggest an essential role for cerebral cortex across species in the perception of these sounds (i.e., gerbils: Wetzel et al., 2008; macaques: Heffner and Heffner, 1984; Harrington et al., 2001; humans: Altman and Gaese 2014; Martin et al., 2024). Cortical encoding of FM sweep trajectories has received significant research attention in multiple models (rat: Zhang et al., 2003; macaque monkey: Tian and Rauschecker, 2004; owl monkey: Atencio et al., 2007; Malone et al., 2017) but essential questions remain, including the roles of spike rate and spike timing as a function of species, behavioral and anesthetic state, and cortical area. Previous studies concentrated their efforts on quantitative analyses of firing rate over fixed intervals relative to stimulus onset (Mendelson and Cynader, 1985; Heil et al., 1992a,b; Eggermont, 1994; Heil and Irvine, 1998; Tian and Rauschecker, 2004), or dynamic intervals defined by the responses themselves (e.g., Atencio et al., 2007), while others captured spike trains via binning techniques (e.g., Malone et al., 2017).

FM sweep trajectories are expected to interact with the frequency areas of auditory neurons, defined as the joint representation of a neuron’s sensitivity to spectral frequency and sound intensity. For example, a neuron tuned to low frequencies would be expected to respond to FM sweeps when a sweep of sufficient intensity passes through that neuron’s best frequency, provided the sweep’s velocity allows sufficient time to integrate acoustic energy in that neuron’s passband. For slower sweeps, the neuron’s spectral bandwidth could be expected to determine the duration of the neural response (e.g., Atencio et al., 2007; Godey et al., 2005). This has important implications for FM sweeps with varying velocities and fixed endpoints, such as those employed in the current study, since FM sweep trajectories affect the expected timing of the neural response, and thus the discriminability of such responses. For example, a neuron tuned to low frequencies would be expected to respond to descending, logarithmic FM sweeps of varying velocities at staggered latencies relative to stimulus onset. However, for ascending sweeps, that neuron’s responses would likely overlap substantially. Our analyses in the current report confront this concern very directly (see below).

Although cortical responses to FM sweeps have been widely studied in multiple species (see above), very few studies have been done in non-human primates (NHPs) (e.g., Malone et al. 2017) and these have been largely restricted to anesthetized animals (Tian and Rauschecker, 2004; Godey et al., 2005; Kajikawa et al., 2008) and/or primary auditory cortex (Atencio et al., 2007). Although there is little data for alert NHPs, imaging studies in human subjects have shown activations throughout auditory cortex to such stimuli (Altmann and Gaese, 2014; Joanisse and DeSouza, 2014; Okamoto and Kakigi, 2015; Martin et al., 2024).

An often cited hypothesis in the literature on primate auditory cortical processing is that there are at least two processing streams, a dorsal ‘spatial’ stream and a ventral ‘non-spatial’ stream (Rauschecker 1998; Rauschecker and Tian, 2000). This is due to first, the local cortico-cortical connections between core auditory areas such as A1 and the rostral field (R), and their immediate neighbors making up the ‘belt auditory cortical fields, including the middle medial field (MM), potential ‘caudal / spatial’ processing fields the caudomedial field (CM) and caudolateral field (CL), and the more ‘rostral / non-spatial’ processing area, the middle lateral field (ML). Secondly, the ultimate pathways project to parietal and frontal lobe areas consistent with similar projections in visual cortex (see Romanski et al., 1999a,b; Rauschecker and Afsahi, 2023). Comparisons of the physiological properties of neurons to briefly presented tone stimuli in core and belt cortical fields, based on traditional metrics of spectral tuning, intensity tuning, and spike latency, have shown that they are distinctive (Merzenich and Brugge 1993; Recanzone, 2000a; Rauschecker and Tian, 2004; Kusmierek and Rauschecker, 2009). When investigating more complex acoustic stimuli, there is compelling evidence that spatial tuning is enhanced from the core (A1) and belt (CL) cortical areas (Recanzone et al., 2000b; Woods et al., 2006; Miller and Recanzone, 2009; Engle and Recanzone, 2013). However, there is less evidence for non-spatial preferences across hierarchical cortical areas (spectral tuning: Ramamurthy and Recanzone, 2016; intensity tuning: Ramamurthy and Recanzone 2020). Those studies compared non-spatial stimulus parameters between the core area A1and belt area CL, which is hypothesized to be more selective for spatial parameters and strong differences were not observed. However, when coding schemes go beyond simple firing rates, differences have been noted for more temporally complex stimuli such as tone or noise pip sequences (Ng and Recanzone, 2018) or amplitude modulated sounds (Niwa et al., 2015; Mohn et al, 2021; Johnson et al., 2020, 2025).

In the present study, we sought to compare the responses of single auditory cortical neurons to logarithmic FM sweeps in alert macaque monkeys across cortical areas in both core (A1 and R) and belt areas (MM, CM, CL and ML). If there is selective processing of FM sweeps in the cortical hierarchy one would predict that the more rostral fields (R and ML) would be more selective to these stimuli than more caudal fields (CM and CL). However, sampling bias could artificially lead to erroneous conclusions. For example, recording chambers that are more caudally located will necessarily oversample low frequency selective neurons in the rostral areas, biasing the sample toward preferring upward sweeps. To eliminate this concern we took considerable pains to ensure that sampling bias could not account for our results, and compared responses of neurons in all regions to those of primary auditory cortex (A1). These results are the first systematic study of responses to non-spatial, time-varying stimuli across core and belt cortical areas in the alert macaque monkey.

## Methods

This study presents novel analyses of data collected during previous studies from our laboratory (Recanzone, 2000a; Woods et al., 2006; Juarez-Salinas et al., 2010; Engle and Recanzone, 2013; Ramamurthy and Recanzone, 2016, 2020), and full details regarding data collection procedures can be found in those publications. The experimental group consists of data from three adult male rhesus macaque monkeys (F, G, and L), aged 5.1-11.5 years during the course of these studies. All procedures used in these experiments were in accordance with AAALAC and the NIH guidelines for the use of animals in research and were approved by the Institutional Animal Care and Use Committee of the University of California, Davis.

### Acoustic stimulus presentation

Experiments were performed in a double-walled anechoic chamber lined with sound-absorbent foam. The monkey was seated in an acoustically transparent primate chair during data collection. Stimuli were comprised of frequency modulated tones (5 ms rise/fall) generated using a Tucker-Davis Technologies (Alachua, FL) system, presented from a speaker located directly opposite to the contralateral ear. Intensities of stimuli tested were 65 +/- 2.5 dB SPL.

### Recording procedures

Monkeys were implanted with a restraining head post and a recording chamber to facilitate orthogonal penetration of the superior surface of the superior temporal gyrus (see Recanzone, 2000a, Overton et al., 2017, for further details) in the left hemisphere. A tungsten microelectrode (2-4 MOhms, FHC Inc., Bowdoin, ME) was inserted through a guide tube (penetrating the brain by ∼3mm), and then lowered using a hydraulic microdrive (Narshige, Japan) during recording sessions. An experimenter seated outside the sound isolation chamber monitored neural activity on an oscilloscope and through an audio speaker while search stimuli (broadband noise bursts, tones, band-passed noise, clicks) were presented. When stimulus driven activity was detected, single units were isolated using a time-amplitude window discriminator (Bak Electronics Inc., Umatilla, FL) and time-stamps of isolated units were recorded with 1 ms resolution. The position of each recording site within core or belt areas was assigned based on location within the recording cylinder and physiological characteristics including tone frequency bandwidth, latency, spatial tuning, and spectral bandwidth selectivity (see Recanzone, 2000a). Electrode track locations for recording sites were verified *post hoc* by histologic analysis and confirmed the physiologically-defined cortical areas.

### Behavioral task

Monkeys performed a simple task during data collection to ensure alertness throughout the recording session. Trials were initiated by depressing a lever, and an acoustic stimulus (S1) was presented which could consist of FM sweeps as described in this report, or other stimuli used for different studies (i.e. broadband noise, tones, etc.; see below). After three to eight more stimuli with different parameters (intensity, tone frequency, spatial location, etc.) were presented (ISI of 750-800 ms), the test stimulus (S2) was presented from a different spatial location (see Ng and Recanzone, 2018). Monkeys were required to release the lever within 800 s following S2 offset to receive a fluid reward, followed by a brief delay to allow time for swallowing before initiation of the next trial. If the monkey did not release the lever in the response window, no reward was provided and instead a brief time-out occurred.

Table 1 shows the numbers of neurons subjected to these FM sweep stimuli in the three monkeys in the different core and belt cortical areas. As described below (**Creation of the Data Sample**), we used a series of criteria to subject the cells to further analysis. Neurons that reached these criteria are shown as the numerator and the total analyzed is shown in the denominator within the columns of Table 2. We also recorded a number of neurons from bordering auditory cortical fields (i.e. AL, RTL, RT, RTM, see Hackett et al., 1998) as well as some from non-auditory cortex outside the belt region that allowed us to physiologically characterize the different cortical fields. The numbers of such neurons within these different cortical areas were too small to make meaningful comparisons so were omitted from further analysis.

**Table 1.** Numbers of neurons recorded from for this experiment. The total of numbers of neurons that were significantly responsive (numerator) is shown with respect to the total of neurons encountered (denominator) for each monkey (L, G and F; columns 2, 3 and 4) for each cortical field (column 1). The vast majority of encountered neurons were responsive by these criteria. The rest of this report is based on those neurons (n = 1090).

| Field | Monkey L | Monkey G | Monkey F | Total | % Responsive |
| --- | --- | --- | --- | --- | --- |
| A1 | 118 / 132 | 194 / 216 | 49 / 51 | 361 / 399 | 90% |
| CL | 139 / 139 | 0 / 0 | 34 / 43 | 173 / 182 | 95% |
| CM | 72 / 86 | 98 / 116 | 34 / 47 | 204 / 249 | 82% |
| ML | 64 / 75 | 81 / 100 | 11 / 11 | 156 / 186 | 84% |
| MM | 39 / 41 | 0 / 2 | 13 / 17 | 52 / 60 | 87% |
| R | 0 / 5 | 67 / 76 | 77 / 79 | 144 / 160 | 90% |
| Sum | 432 / 478 | 440 / 510 | 218 / 248 | 1090 / 1236 | 88% |

**Table 2.**
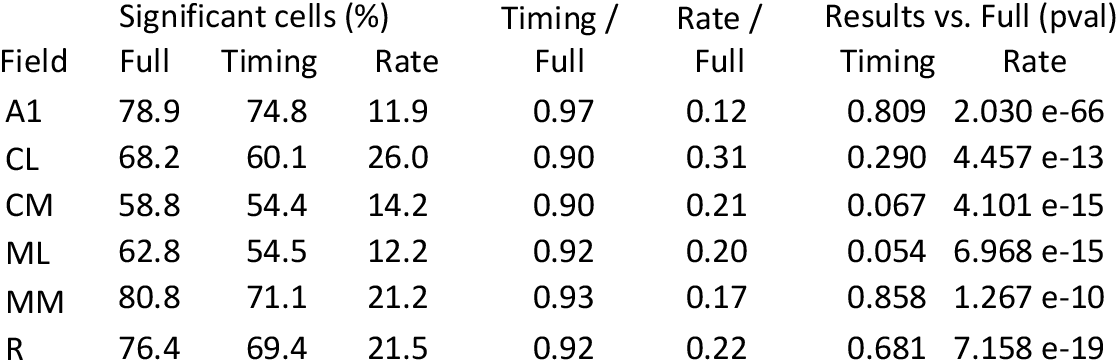
Comparison of significant cells by cortical area and decoder. The first set of columns show the percentage of neurons that showed significant decoding accuracy (above the heavy dashed line of Fig. 5) using the full spike train (‘Full’), the timing only decoder (‘Timing’) or just the overall firing rate (‘Rate’). The middle two columns show the ratio of the mean decoding values above chance relative to the full spike train. p-values are shown in the last two columns.

### Calculation of BF

We measured frequency tuning to static pure tones by presenting 14 frequencies spanning the majority of the FM sweep range (0.5, 1, 2, 3, 4, 5, 6, 7, 8, 9, 12, 14, 16, 18 kHz) at 65 +/- 2 dB SPL. Tones were presented 12 times in pseudorandom order, interleaved with other stimuli including the FM sweeps. We determined the spike count within 100 ms of tone onset for all cells, and defined the nominal best frequency as the tone frequency that produced the largest spike count (resolving ties as the lowest tone frequency among the list). In a subsequent block of trials, the frequency eliciting the best response as defined above was presented for 12 trials at four different intensities (45 – 75 dB SPL) in 10 dB increments.

### FM Sweeps

We presented a set of 10 logarithmic frequency modulated (FM) sweeps spanning a range of 500 Hz to 20 kHz (5.32 octaves) at 5 distinct velocities and 2 directions (‘up’: 0.5 – 20 kHz and ‘down’: 20 – 0.5 kHz). The durations of the sweeps were 60, 120, 180, 240, and 350 ms, corresponding to velocities of 71.4, 35.7, 23.8, 17.9, 12.2 octaves/second. Sweeps were presented in pseudorandom order; each sweep was presented 12 times. For convenience, upsweeps are defined as having positive velocities and downsweeps as having negative velocities. Figure 1 shows the frequency (kHz) as a function of time for both types of sweeps. Horizontal lines show the time duration from sweep onset to each of the 13 frequencies (1 – 18 kHz) for the five different sweep durations in both the upward (Fig. 1A) and downward (Fig. 1B) directions. These physical parameters of the stimulus, while consistent across octave space, nonetheless show how simple timing codes depending on the best frequency of the neuron could trivially, and in principle, be used to decode both the direction and speed of these stimuli, which is a major consideration for the resulting analysis. For example, a neuron with a best frequency of 9 kHz would have distinctive responses in time relative to those of a neuron with either a 1 kHz or an 18 kHz best frequency depending on the direction and speed of the FM sweep.

**Figure 1.**
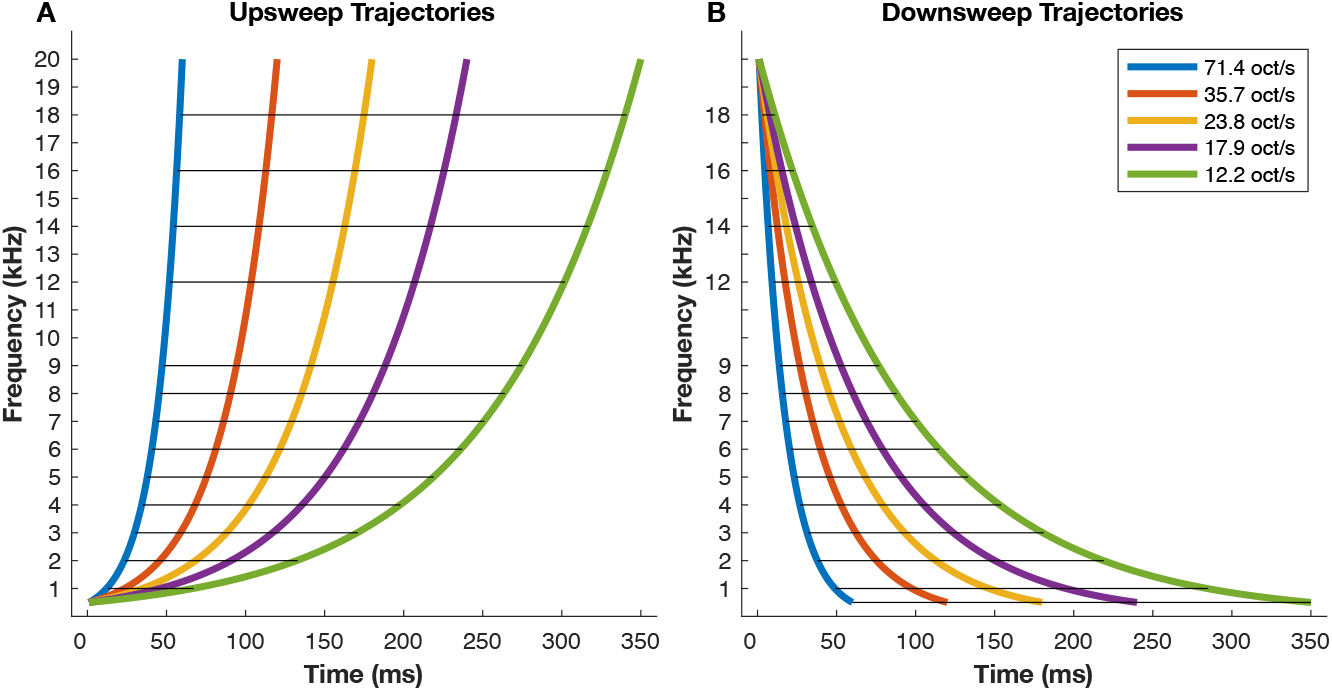
FM sweep stimuli used in this experiment. Sweeps were presented at 5 velocities (colored lines, see inset), in upward (A) and downward (B) directions. Each sweep spanned a frequency range from 0.5 to 20 kHz. The horizontal black lines illustrate the intersection latencies for the frequencies that were presented as tones to determine the best frequency for each cell (excepting 0.5 kHz). As is evident, the span of the intersection latencies, indicated by the length of each line, vary widely for sweeps in different directions (up or down) at the extremes of the frequency range.

### Spike Train Decoding

As in prior work, we use the term “decoder” to refer to the algorithms used to estimate stimulus density from neural responses (Malone et al. 2017). Spike trains were classified on a trial-by-trial basis by a nearest neighbor linear decoder that computes the Euclidean distance between each spike train and the mean PSTH for each stimulus in the set. The stimulus corresponding to the mean PSTH ‘nearest’ the spike train is estimated to have elicited that spike train. The spike train from the trial being decoded is excluded from calculation of the mean PSTH for that stimulus via ‘leave-one-out’ cross validation. The distance between two spiking responses is defined in a metric space based on binned spike counts, such that each time bin represents a dimension in the space and the spike count for each bin represents the coordinate along the corresponding dimension. Decoding accuracy was reported as the percent of total trials that were correctly assigned to the eliciting stimulus. Optimal bin size was defined as the bin size that maximized decoding accuracy. Because mutual information increases monotonically with increases in decoding accuracy (Schnupp et al., 2006; Malone et al., 2007, 2017) we sometimes describe neurons with high decoding accuracy as “informative”.

Because the duration of the analysis interval was 400 ms (see **Peak Decoding**, below), binning the responses at 400 ms (i.e., in a single bin) results in a decoder that relies entirely on average firing rate information, which we term the ‘rate-only’ classifier. It is also possible to remove differences in average firing rates across stimuli while retaining information about differences in how spikes are distributed in time by normalizing each test and each template by the spike count. In prior work, we had normalized each test and template by its respective vector norm, which entails that responses to different stimuli are mapped to an equivalent distance from the origin in the response space. Both normalization procedures produce extremely similar results (data not shown), but spike count normalization is arguably simpler. We refer to this as the timing-only classifier.

When decoding the frequency of tone pips, we computed the “error cost” (Malone et al., 2010), which assigns a cost proportional to the ordinal difference between the actual stimulus and the estimate by computing the distance of each estimate from the correct entry in the confusion matrix (i.e., the within-column distance to the diagonal). For example, assigning a label of 7 kHz for a tone at 8 kHz would incur a cost of 1, while assigning a label of 6 kHz would incur a cost of 2. Correct assignments incur a cost of 0. The error cost represents the sum of costs across all stimuli and trials.

### Calculating Significance for Spike Train Decoding

We assigned significance values for decoding accuracy by comparing actual confusion matrices to simulated confusion matrices of matching size populated by random draws from a uniform distribution spanning the number of elements in the stimulus set (e.g., 10 for all FM sweeps, 5 for FM sweeps in a cell’s better direction, etc.). The number of random draws for each stimulus was equal to the number of experimental trials, so each column of the simulated confusion matrix summed to the number of repeated trials for each unique stimulus. The decoding accuracy for each genuine confusion matrix was compared against the distribution of simulated decoding accuracies, and p-values were assigned by counting the fraction of simulated accuracies that exceeded the actual accuracy, divided by the total number of simulated values (n = 1,000,000). We assigned significance values for error cost analogously.

### Calculation of the optimal binwidth

We computed decoding accuracy for all sweeps using the following binwidths (in ms): 1, 2.5, 5, 10, 15, 20, 25, 50, 100, and 400; We measured the normalized mean decoding accuracy, which was maximum at 20 ms (86.3%), indicating that this was nearly optimal for a large number of cells. For every cortical field, 20 ms was the modal optimal binwidth, with the exception of field CM, where the modal optimal binwidth was 15 ms (20% of cases), and the second most common optimal binwidth was 20 ms (17% of cases). Based on these results, we used 20 ms bin size for the timing-only classifier, and 400 ms for the rate-only classifier for all analyses to follow.

### Redressing BF imbalances

In order to minimize the impact of mismatches in the BF across cortical fields, we employed a greedy matching algorithm that stepped through each cell from a non-A1 field and identified an A1 cell that minimized the absolute value of the difference in both cells’ BF. Since A1 was oversampled relative to the other fields (see Table 1), there were usually multiple matches for each non-A1 unit. We chose the first match, then removed the matching cell from the list of potential matches (i.e., matches occurred without replacement). To make our comparisons conservative, we only included non-A1 cells where an exact BF match could be identified in A1. This represented 100% of cells in CL, 77% of cells in CM, 98% of cells in ML, 100% of cells in MM, and 100% of cells in R. These BF-matched subpopulations were then used to verify that differences in decoding accuracy were unlikely to be derived from sampling differences related to frequency tuning among the different cortical fields.

### Peak Rate Decoding

We evaluated whether identification of a particular response window could improve the performance of the rate-only classifier. To do so, we identified the portion of each PSTH that maximized the spike count for a fixed window size, and fed the trial-by-trial spike count within this window to the rate-only classifier. We tested a range of analysis window intervals from 25 to 500 ms in 25 ms intervals. A window size of 400 ms provided the highest accuracy across the sample, demonstrating that even optimizing firing rate information on a cell-by-basis by precise windowing of the responses does not improve the performance of the rate-only classifier. Based on this analysis, the interval from 0-400 ms was used as the analysis window for all spike train classifiers.

### Direction Selectivity Index (DSI)

We computed the DSI by summing spike counts across trials for matched-velocity sweeps on the interval from 0-400 ms. The DSI was computed as the ratio of the difference in spikecounts (upsweeps-downsweeps) to the sum of spikecounts for sweeps in both directions. By this metric, responses that were twice as strong to upsweeps than to downsweeps would yield a DSI of ∼0.33 (e.g., (2-1)/(2+1)). DSI values were calculated separately for each sweep velocity, and each cell’s responses were captured as a vector of 5 DSI values. To determine whether a given neuron exhibited a consistent preference for sweep direction across different sweep velocities, we relied on the assumption that a genuine preference for sweep direction would be consistent across sweep speeds, and thereby reduce the variance in DSI values. We computed the variance of the DSI values across the 5 tested sweep speeds for each neuron in the sample. We estimated a null distribution of DSI variance by randomly selecting 5 DSI values (one from each data column representing velocity) and computing the variance for a simulated “cell” (n = 100,000). We assigned p-values based on the number of times that the simulated DSI variance was smaller than the actual DSI variance.

### General Statistical Methods

Our null hypothesis for our statistical tests was that non-A1 fields did not differ from A1. We organized the analyses to determine whether neural representations of FM sweeps had changed significantly with respect to A1 to limit the number of statistical comparisons, and because this provided a simple conceptual framework for understanding our results. We report uncorrected p-values for all statistical tests exactly, so readers can draw their own conclusions (Heroux et al., 2016). When significance tests were used for inclusion in the database or in specific analyses, we avoided excluding cells wherever possible to limit reducing the statistical power of tests based on comparing distributions of values. When comparing distributions, we employed Kolmogorov-Smirnov two sample tests (MATLAB kstest2) to test whether data from different cortical field came from the same continuous distribution.

### Creation of the Data Sample

Our inclusion procedure was intended to be maximally inclusive while excluding nonresponsive neurons. We were concerned that rigorous exclusion of marginally informative neurons might obscure differences among the cortical fields. This procedure involved the union of 3 tests: 1) Did stimulus driven activity exceed the spontaneous rate during the initial 100 ms of the tonal stimuli used to determine the frequency tuning function (p < 0.01)? 2) Did a cell exhibit significant tuning to tone pips, such that the response at the best frequency exceeded that to the worst frequency (p < 0.05), or successfully discriminated among various tones? 3) Did a cell exhibit decoding accuracy above chance for the complete set of FM sweeps? Any cell that produced a significant result on any of these criteria was included in the database, resulting in 1090 out of 1236 cells that were unambiguously localized to one of the six cortical areas (see Table 1).

For the first criterion, we compared spike counts during the spontaneous interval (350-450 ms prior to trial onset) to responses to tones during the interval from 25-125 ms. To make the test more conservative, we calculated the maximum spontaneous rate across tested frequencies after averaging across trial repetitions (n = 12) to compare against the best frequency (BF). We compared spike counts within the analysis window with a two-sided Wilcoxon sign rank test (Matlab signrank), and included cells where the p-value was less than 0.05. The second criterion was intended to capture neurons that responded at significantly different rates to different tone frequencies. We compared the spike counts from all trials for the best frequency and worst frequency, using the same test and criterion. For the third criterion, we also applied the full spike train classifier to the tone responses, and computed the error cost from the resulting confusion matrices (see *Spike Train Decoding*, above) to capture additional information about tone frequency encoding. Cells that exhibited smaller error costs than expected by chance were included in the database. We also assigned significance values to the decoding accuracy of the full spike train classifier for all sweeps and included cells where performance exceeded our significance criterion (p < 0.01).

## Results

This work seeks to determine whether auditory cortical fields outside of primary auditory cortex (A1) differ in terms of the quality or nature of their encoding of complex, dynamic acoustic signals. However, the ability to decode exponential sweeps on the basis of direction and velocity is partially confounded by how the sweeps interacts with cortical frequency tuning. The lengths of the horizontal black lines on Figure 1 indicate the difference in the latency at which sweeps at different velocities intercept the frequencies depicted on the ordinate of the graph. We plotted the lengths of those lines as a function of frequency in Figure 2. These plots show that neurons tuned to frequencies near 3 kHz would be expected to differentiate FM sweeps in both directions effectively if an important component of that differentiation is the latency of the neural response relative to stimulus onset (Godey et al., 2005; Atencio et al., 2007) and that the neuron has knowledge of when the sweep onset occurred. Since all sweeps originate and terminate at fixed endpoint values (0.5 and 20 kHz), there is a considerable difference in the latency span at different frequencies. For example, upsweeps originating at 0.5 kHz will arrive at 1 kHz relatively quickly, regardless of the sweep velocity. By contrast, since the sweep terminus is 20 kHz, upsweeps arrive at 18 kHz at widely varying latencies, approaching the complete sweep durations (see Methods). Because the latency spans are significantly different for different frequencies, we expect that neurons tuned to different frequencies (e.g., 1 kHz or 18 kHz) are likely to exhibit proportionate differences in how their responses are distributed in latency, which in turn affects our ability to decode differences in sweep velocity based on those responses. Similarly, we expect neurons with best frequencies in the middle frequency range to be better able to encode sweep duration and velocity given their expected response latency distributions.

**Figure 2.**
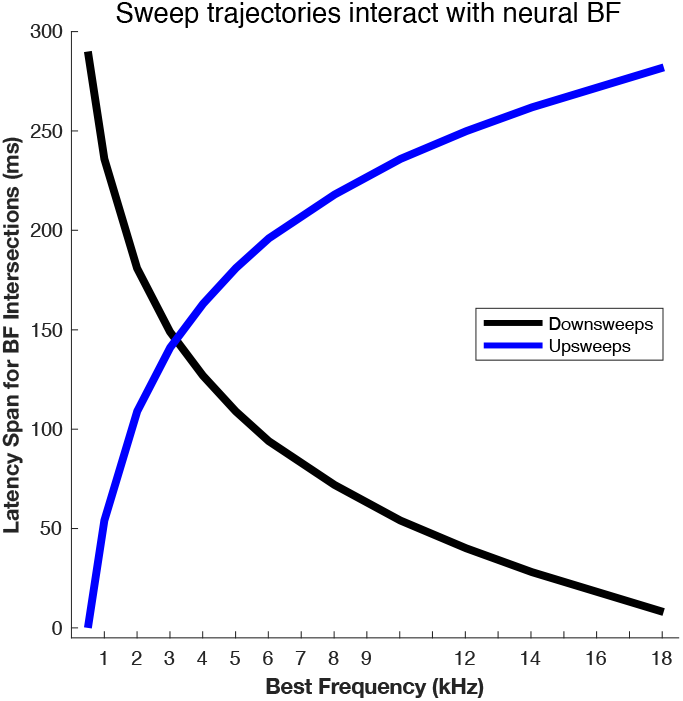
Stimulus latency as a function of frequency relative to sweep onset. Sweeps of different velocities intersect the frequency values mapped to the abscissa, which we refer to as the latency “span”. These differences correspond to the lengths of the horizontal black lines shown in Fig. 1.

If a neuron fires in response to the sweep entering or approaching its response area, that neuron’s best frequency (BF) will determine how widely spaced the responses to sweeps of given velocity will be in the PSTH. Figure 3 shows an example of BF-dependent sweep direction encoding for a neuron with a BF of 18 kHz. The PSTH peaks were clearly displaced in time depending on the sweep velocity for upward sweeps (Fig. 3A). In contrast, responses to downward sweeps (Fig. 3B) are nearly identical regardless of sweep speed. As a consequence, this neuron showed much higher decoding accuracy for upsweeps (96.7%; Fig. 3C) than for downsweeps (28.3%; Fig. 3D) when sweeps in different directions were decoded separately.

**Figure 3.**
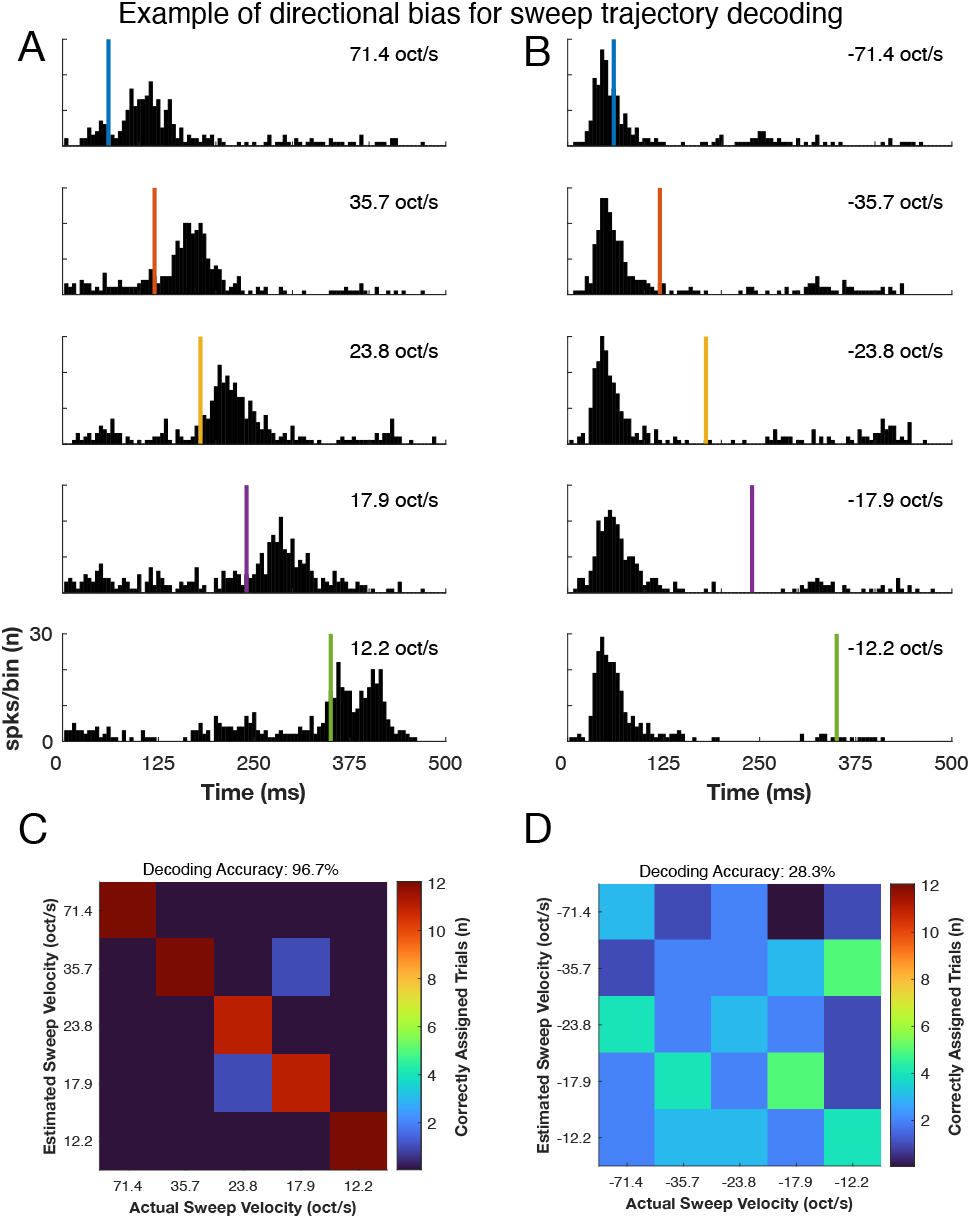
Representative neuron with a high best frequency. A) PSTHs from a cell in ML with a best frequency of 18 kHz showing the binned spikes counts elicited by ten FM sweeps in the upward (left column) or downward (right column) direction. Insets indicate the sweep velocity from fastest (top row) to slowest (bottom row). Vertical red bars indicate the termination of the sweep. Note the clear latency differences for the upward sweeps that is not present for the downward sweeps. B and C) Confusion matrices illustrating the number of correctly assigned trials for the upward and downward sweeps respectively. Values along the diagonal indicate correct assignment.

Figure 4 shows the results for a different cell, recorded in field CM with a BF of 4 kHz. The PSTHs illustrate consistent increases in response latency for slower sweeps for both upsweeps (Fig. 4A) and downsweeps (Fig. 4B). As a result, there is greater parity in the decoding accuracy for sweeps in either direction: 63.3% for upsweeps (Fig. 4C) versus 78.3% for downsweeps (Fig 4D) compared to the cell whose responses were depicted in Figure 3. Nonetheless, this cell does show a decoding accuracy advantage for downward sweeps in the direction predicted by the latency span curves in Figure 2.

**Figure 4.**
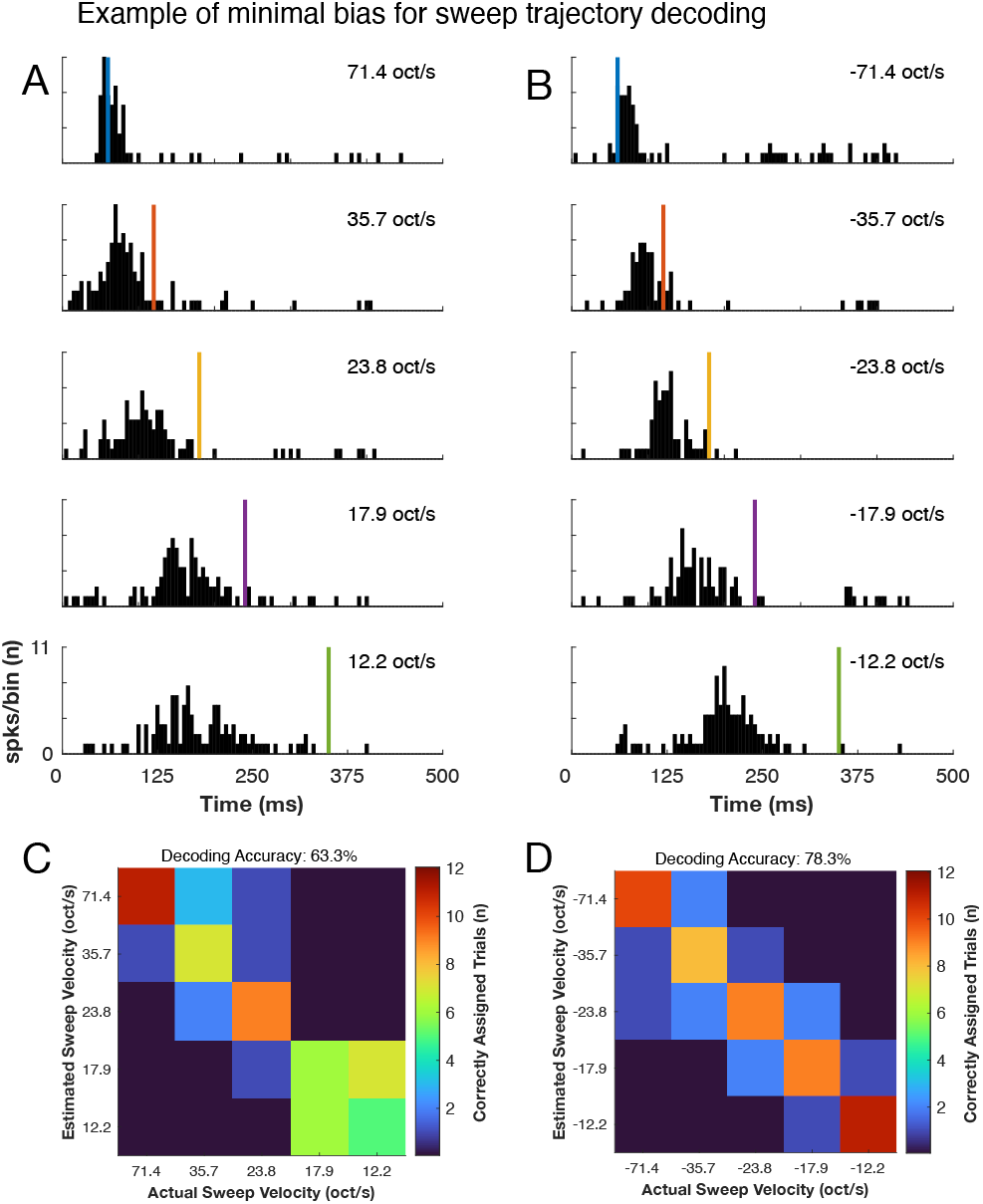
Representative neuron with a middle best frequency. PSTHs and confusion matrices for a cell from CM with a best frequency of 4 kHz. Here there are clear latency differences in the responses for both upward and downward sweeps, With correspondingly better performance of the decoder. Conventions are identical to those in Figure 3.

The examples in Figures 3 and 4 demonstrate the need for careful consideration of how the distribution of BFs might affect decoding accuracy when discriminating FM sweep velocity and direction. We quantified the effect of BF on decoding accuracy at the population level by analyzing correlations between decoding accuracy and neuronal BF. We limited this analysis to cells in the sample that exhibited decoding accuracy significantly above chance at the p < 0.001 level (n = 660). We used a strict criterion here because the inclusion of uninformative neurons simply adds noise to the analysis. Decoding accuracy for upsweeps was significantly and positively correlated with neuronal BF (r = 0.31; p = 9.4464e-16); decoding accuracy for downsweeps was significantly and negatively correlated with BF (r = -0.41; p = 2.6869e-28). The significance of these relationships demonstrates the need for correcting for differences in the distributions of frequency tuning among different cortical fields when comparing decoding accuracy values.

### Correlations between BF and Decoding Accuracy

We visualized the relationships between decoding accuracy, cortical field, and neuronal BF by creating the swarmchart depicted in Figure 5. Each neuron in the database is plotted twice, to illustrate the decoding accuracy, computed separately, for downsweeps (left side of pair) and upsweeps (right side of pair), as indicated by the arrows shown above the data for A1. The color of each dot indicates the neuronal BF (color bar at right), and accuracy values (y-axis) reflect use of the full spike train (FST) classifier. Examination of the A1 data (leftmost pair of swarmcharts) reveals that the highest decoding accuracies for downsweeps (left) were observed for neurons tuned to lower BFs, as indicated by the prevalence of cooler colors for higher decoding accuracies. By contrast, many of the highest decoding accuracies for upsweeps (right) were observed for neurons tuned to higher BFs, as indicated by warmer colors (e.g., orange, red) that were rarely observed for downsweeps at similar decoding accuracies. This trend is easiest to see for A1 neurons, given the larger sample size (see Table 1), but this pattern held for the other cortical fields. Moreover, comparing the colors across different cortical fields also reveals that the distributions of BF varied. There is a relative preponderance of blue dots for field R, for example, and a relative preponderance red dots for field CM, most certainly due to the sampling bias of those fields—low frequencies border A1 in area R, and high frequencies border A1 for field CM (Brugge and Merzenich, 1973; Recanzone et al., 2000a). A1 is much larger in size than the surrounding belt fields (Kaas and Hackett, 2000). This fact, combined with morphological constraints of the macaque skull and recording chamber technology at the time of these studies, limited the region of cortical space we could investigate (Recanzone et al., 2000a).

**Figure 5.**
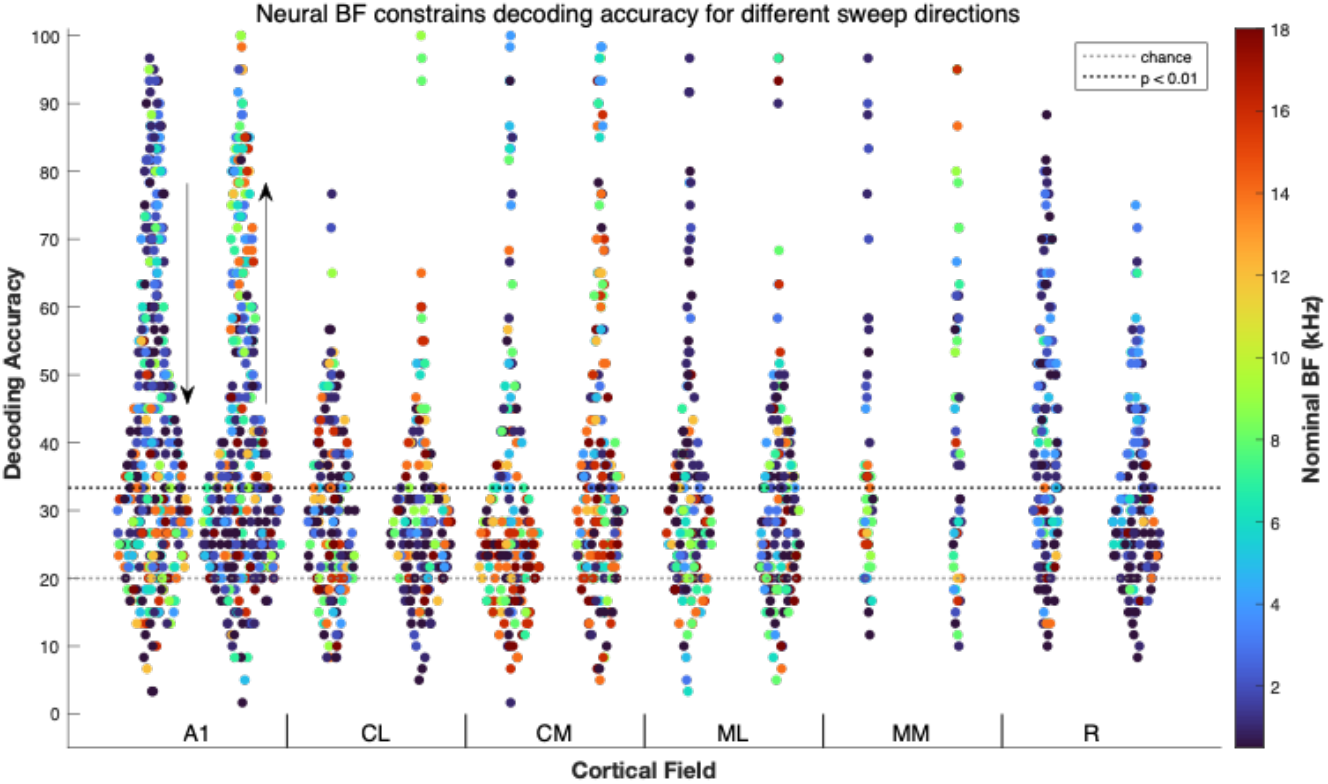
Decoding accuracy for sweep direction using the entire spike train. Swarmchart plots showing the decoding accuracy for downsweeps (left grouping) and upsweeps (right grouping) for neurons (n = 1090) recorded from the labeled cortical fields(See Table 1). Each neuron is thus represented by two points. The dotted lines indicate the accuracy values expected for chance (light grey) and corresponding to a p-value of 0.01 (heavier line) based on simulated confusion matrices (see Methods). Colors for each dot indicate the BF estimated from responses to tone pips. There is very little difference in the distributions of BF or the relative distributions of decoding accuracy across cortical fields.

When we quantified the differences in frequency tuning across cortical fields by computing the mean/median BF values for each of the cortical fields in our data sample, we found no difference in the mean/median values for A1 (3/5.51) versus CL (8/9.39), ML (3/5.66) and MM (4.5/5.35; p > 0.1 for each comparison). As expected, we did see a difference for A1 versus CM (3/5.66) and R (2/3.12). CM exhibited higher BFs (p = 1.316e-10) and field R exhibited lower BFs (p = 7.396e-06; MATLAB kstest2) relative to A1.

The overrepresentation of high BFs in CM and low BFs in R manifested as significant differences in the decoding accuracy for sweeps in different directions. In CM, median accuracy was 31.7% for upsweeps, and 25.0% for downsweeps. In R, median accuracy was 28.3% for upsweeps and 39.2% for downsweeps. In the other cortical fields, the medians for upsweeps/downsweeps did not differ significantly (A1: 33.3%/35.0%; CL: 28.3%/30.0%; ML: 29.2%/28.3%; MM: 36.7%/31.7%; p > 0.1 in all cases). This pattern of results is consistent with the notion that sweep velocity is primarily decoded by spike timing differences related to the latency of the intersection of the sweep and each neuron’s BF, since the interaction of the sweep trajectory affects when spikes occur, but not necessarily how many spikes occur.

This pattern of results also demonstrates the need for care when making comparisons across cortical fields. Figure 5 suggests that decoding accuracy was superior in A1 relative to nonprimary fields other than MM, which was under-sampled relative to the other fields due to its relatively small size and the difficulty of recording in that field in this preparation (see Kaas and Hackett, 2000; Recanzone, 2000a; Kusmierek and Rauschecker, 2009). Nevertheless, the upper limit on decoding performance among belt neurons was quite similar to that of the core fields, A1 and R, with the possible exception of field CL. It is also clear that the variance in decoding accuracy within each field was substantial. Given this variance, we encountered informative and uninformative neurons in each sampled field.

Although the data in Figure 5 does not indicate large differences in the responses of neurons across cortical fields, further analysis confirmed their significance independent of sampling bias related to BF. We eliminated BF as a confound by using a greedy matching procedure to downsample cells recorded from A1 to attempt to match the BF distributions from the nonprimary fields (see Methods). Given the comparatively large sample of A1 neurons (n = 361), it was possible to identify exact BF matches for all cells in fields CL (n = 173), MM (n = 52), and R (n = 144), 98% of cells in ML (n = 153), and 77% of cells in CM (n = 157). We then compared the distributions (MATLAB kstest2) of decoding accuracy for all sweeps for BF-matched groups against that of AI. Decoding accuracy in A1 significantly exceeded that of all non-A1 fields except MM. Due to the BF matching procedure, median accuracy for A1 changes for each comparison, since different subsets of the full A1 sample are included (see Methods). The results were as follows: A1 versus CL (medians: 27.50%/20.83; p = 4.121e-09); A1 versus CM (25.83%/19.17%; p = 2.512e-04); A1 versus ML (26.67%/19.17%; p = 4.175e-05); A1 versus MM (27.50/26.67; p = 0.99); A1 versus R (26.67/24.17; p = 1.079e-02). Although we are reporting medians here for convenience, the significance values reflect differences in the accuracy distributions, not the medians. Thus, we saw no real evidence of an enhancement of FM encoding from A1 to more rostral or belt fields, indicating that there is no systematic processing of FM stimuli along this dimension of hierarchical processing.

Median decoding accuracies were relatively low in all cortical fields, due to the presence of large numbers of uninformative neurons among the sound responsive neurons in our sample. Among belt fields CL, CM, and ML, the accuracy distribution was centered below significance, but each exhibited long tails extending to relatively high accuracy. Figure 5 suggests that the transition from A1 to belt fields, excepting MM, was marked by a reduction in the prevalence of highly informative neurons, but this did not appear to act as a lower ceiling on the quality of FM sweep encoding. These analyses indicate that there is not an exclusive hierarchy where FM decoding is dramatically reduced in subsequent processing stations, even if neurons in A1 are generally more accurate at decoding FM direction and velocity compared to neurons in other core and belt areas.

Having evaluated decoding accuracy distributions for full spike trains, we turned to the question of whether temporal or rate encoding more accurately reflect higher order cortical area processing. To this end, we also computed decoding accuracy for two additional classifiers, the timing-only and rate-only classifiers (see Methods; Malone et al., 2007), to complement results for the full spike train classifier depicted in Figure 5. Comparison of results for the three classifiers allows us to parse the contributions of spike timing and spike rate to the information present in cortical spike trains. For example, one might predict a shift from a temporal to a rate code from core to belt cortical areas, similar to that seen for vector strength from peripheral to cortical auditory locations (see Joris et al., 2004). According to this model, the performance of rate-only classifiers (i.e. ‘tuned’ neurons) should increase along the cortical hierarchy, at least in the more rostral fields given the proposed parallel pathway organization of the primate auditory cortex (Raushchecker, 1998; Kaas and Hackett 2000; Rauschecker and Afsahi, 2023; Recanzone and Cohen 2009).

Figure 6 depicts the distributions of decoding accuracy (y-axis) based on the full spike train (left), timing-only (middle), and rate-only (right) classifiers for each of the six sampled cortical fields. The color bar (z-axis) shows the BF from low frequency (cool colors) to high frequency (warm colors). Comparison of the rightmost swarmchart of each triad to the others demonstrates that spike rates provide relatively little information about FM sweep trajectories—the full spike train and timing-only distributions are highly similar, while the rate-only distribution exhibits a median generally above but still near chance performance. There are no differences across cortical areas, particularly in ML and R, that suggest a greater rate-based selectivity for FM sweeps.

**Figure 6.**
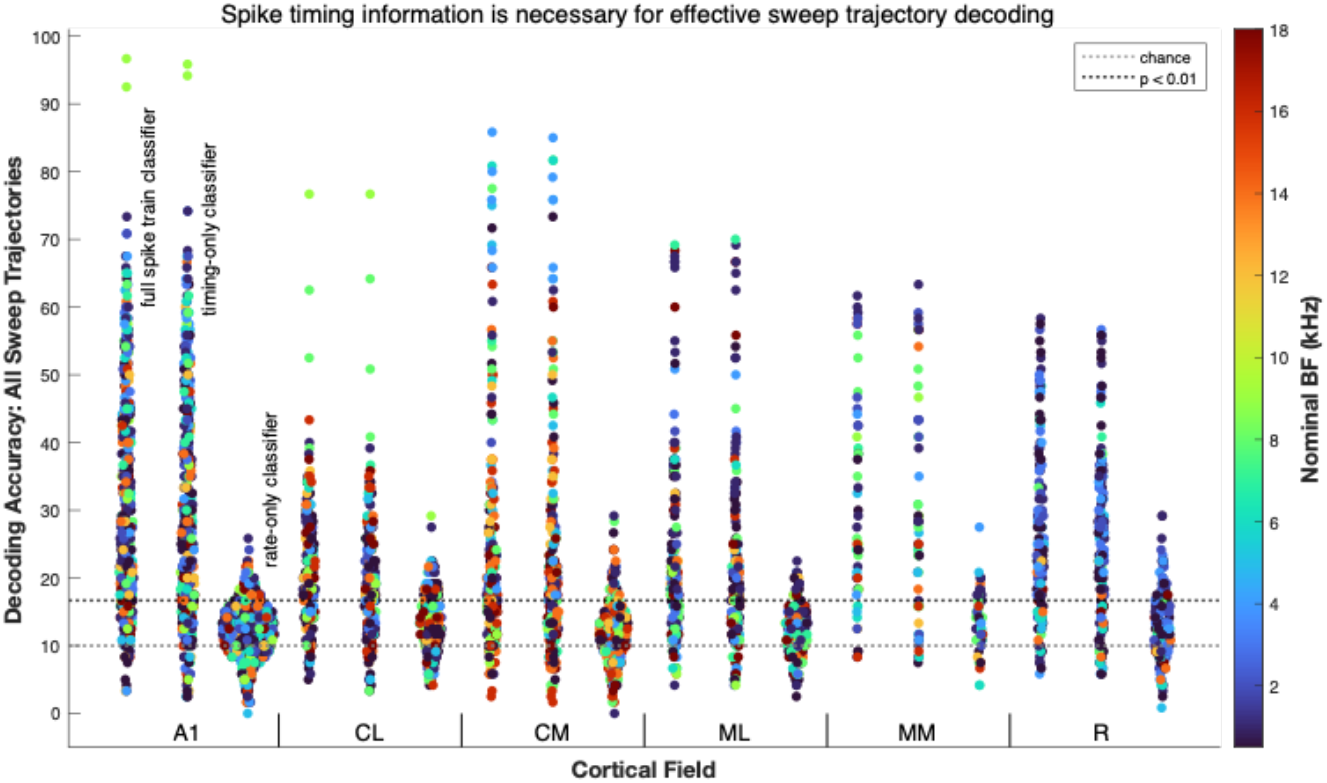
Decoding accuracy for three methods used. Swarmchart plots of the decoding accuracy for all sweeps in three groupings corresponding to the different classifiers (see Methods) labeled above the data for A1. Each neuron in the data sample (n = 1090; see Table 1) contributes three points to this analysis, one for each classifier. The dotted lines indicate the accuracy values expected for chance (light grey) for a set of 10 distinct sweeps and corresponding to a p-value of 0.01 (heavier line) based on simulated confusion matrices (see Methods). Colors for each dot indicate the BF estimated from responses to tone pips. There was very little difference between the full spike train (left most) and timing-only (middle) decoders, but the rate-only decoder (right most) showed much worse classifier performance for neurons in all six cortical areas.

To quantify these impressions, we calculated the percentage of cells that exceeded the significance criterion for each classifier, and for each cortical field, as shown in columns 2-4 in Table 2. The percentage of significant cells was highest in A1 and lowest for ML and CM for both the full and timing-only based classifiers, with the other three fields somewhere in between. The rate-only classifier resulted in far fewer cells that exceeded the significance criterion. A1, CM and ML showed the lowest percentages (<15%).

We quantified differences in encoding quality for different spike train features by calculating the ratio of decoding accuracy for the timing-only and rate-only classifiers to that of the full spike train classifier. Because we were interested in the mutual information between the stimuli and the decoded responses, we subtracted the accuracy expected by chance (i.e., 10% for a set of 10 sweeps) before calculating the ratio, since chance performance is expected when there is no useful information. Cells that did not exhibit significant decoding accuracy for the full spike train classifier were excluded from this analysis. We computed the median ratio for each cortical field to obtain the results shown in the 5th and 6th columns of Table 2. A large majority of information supporting decoding accuracy for sweep trajectory could be extracted from spike timing (90-97%) and only a minority could be captured by spike rate (12-31%).

We did not observe significant differences (kstest2; Matlab) between the full spike train and timing-only classifiers in any tested field (column 7 of Table 2). By contrast, the median differences between the full spike train and rate-only classifier were substantial. The reduction in the median differences for these belt fields is likely a floor effect, since those reductions in median differences relative to A1 (or MM) are largely proportional to the differences in decoding accuracy relative to A1 (or MM). Distributions of decoding accuracy for the timing-only classifier significantly differed from those of the rate-only classifier in all fields, as would be expected from the profound differences evident in Figure 6 (column 8, Table 2).

Overall, these analyses indicate that the temporal information across core and belt auditory cortical areas in the primate provides a substantial amount of information with respect to the temporal dynamics of complex auditory stimuli, with little evidence of a shift from temporal to rate coding for these stimuli from primary to nonprimary areas.

Given the concerns about how decoding accuracy interacts with BF, we also computed decoding for FM sweeps in each direction separately. We expected that this would minimize, if not strictly eliminate, the frequency tuning confound brought about by BF tuning near the low or high end of the sweep frequencies (see Fig. 2). For example, neurons with low BFs could use timing information for downsweeps while neurons with higher BF would be advantaged for upsweeps. We therefore determined the sweep direction (up or down) that resulted in the higher decoding accuracy for each cell, and plotted those accuracy values in

Figure 7. The pattern of results is very similar to that depicted in Figure 6, but the accuracy values, as well as the chance and significance levels, are shifted to higher values, since there are 5 rather than 10 sweeps. There were cells in AI, CL, and CM whose responses could be correctly assigned on every trial (100%).

**Figure 7.**
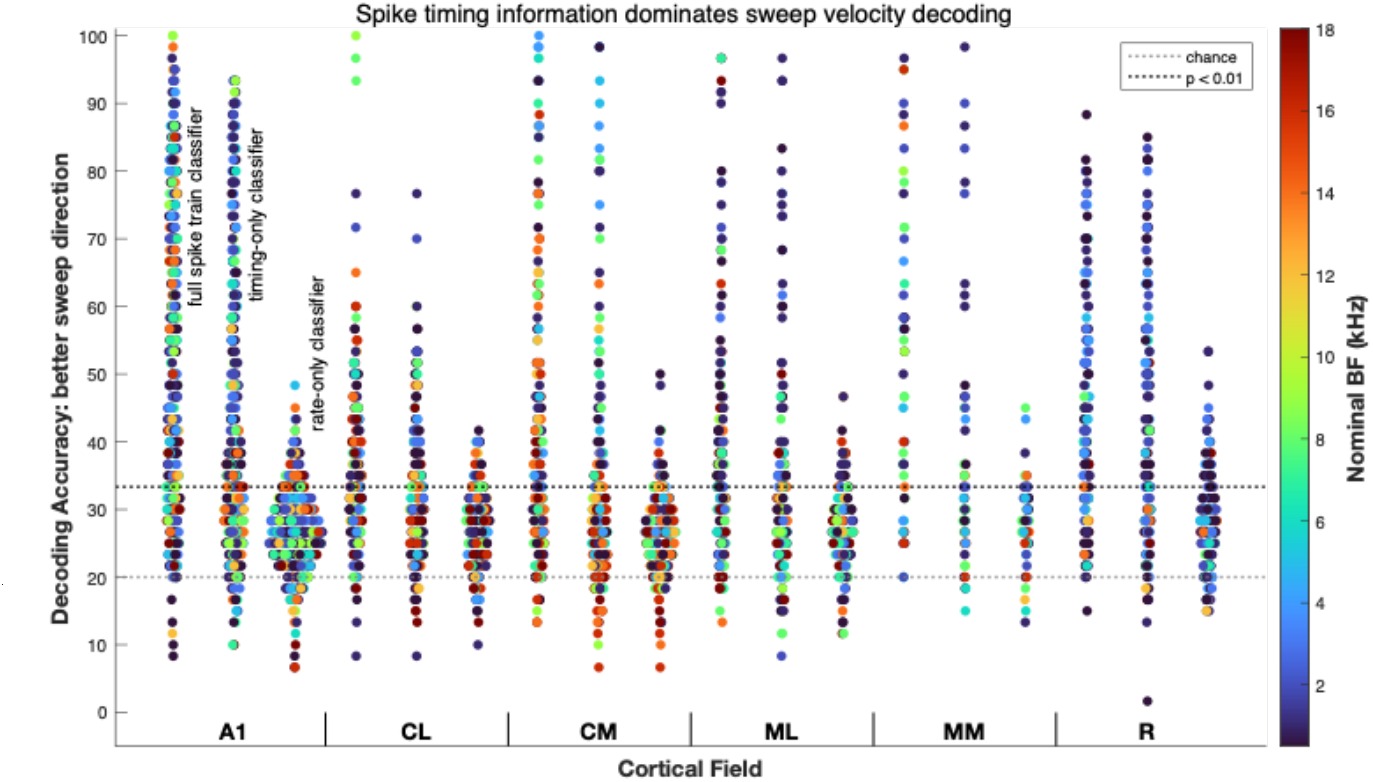
Decoding accuracy for best direction. Swarmcharts for neurons restricted to only the best direction for that cell and are based on data from the five different sweep velocities in that direction. Limiting the analysis to the best direction showed little difference than when both sweep directions were included (compare Figs. 6 and 7) indicating that none of the classifiers were particularly hindered by inclusion of data from cells responding to less-preferred sweep directions. Figure conventions are identical to those in Figure 6.

When we tested for differences among the cortical fields for BF-matched sets, the pattern of results was similar to that for all sweeps, although decoding accuracy in R and MM was slightly higher than that of the BF-matched A1 subsamples, though not significantly so. Results for the better direction for each cell were as follows: A1 versus CL (medians: 46.67% / 33.33%; p = 1.132e-11); A1 versus CM (41.67% / 33.33%; p = 6.462e-04); A1 versus ML (45.00% / 35.00%; p = 1.292e-06); A1 versus MM (44.17 / 31.67; p = 0.99); A1 versus R (41.67 / 42.50; p = 2.273e-02). Here again the p-values represent the outcomes of statistical tests on the distributions (kstest2), not the medians. As we observed when decoding sweeps in both directions, the belt fields CL, CM, and ML exhibited significantly lower median decoding accuracy, reflected in the differences in the decoding accuracy distributions.

We also analyzed the prevalence of direction selectivity, here defined as a rate-based preference for a FM sweep at a given velocity in a particular direction (Figure 8). We operationalized the notion of a “rate-based preference” as significant decoding accuracy for the rate-only classifier applied to sweeps of different directions but the same duration (or velocity, since the frequency span of all sweeps was fixed). Overall, the prevalence of direction selectivity was low across all sweep durations across all cortical fields 60 ms: 5.6%; 120 ms: 6.8%; 180 ms: 6.1%; 240: 7.7%; 350 ms: 8.8%).

**Figure 8.**
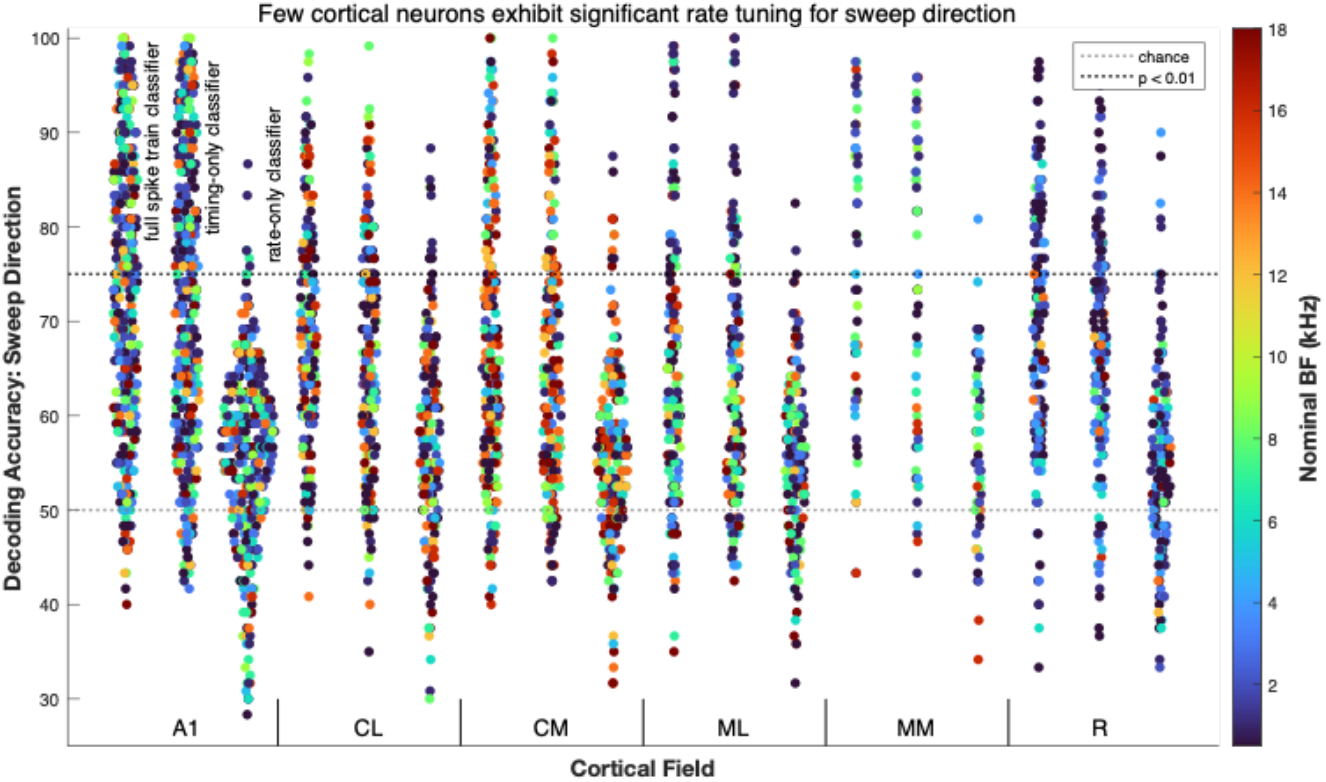
Decoding accuracy for sweep direction across velocities. Sweep decoding was performed for pairs of matched velocities in different directions. Note the increase in the level of significance (heavy dotted line) given that the decoder only compared matched velocities in different directions. Similar trends of the decoding accuracy being much worse under the rate-only condition are still apparent in this case as well. Figure conventions are similar to those in Figure 6 and 7.

This is evident by examining the distributions of decoding accuracy for the rate-only classifiers in Figure 8, where few points in the rightmost swarmchart for each cortical area are significantly above the significance level of p < 0.01 (uncorrected; dark dotted line).

Nevertheless, there is a visible trend where decoding accuracy for the rate-only classifiers was above chance (50%) but below our significance criterion. The implication is that a population-based decoder could leverage that information to decode FM sweep direction more effectively than single-cell based decoders. Of course, population-based decoders that leverage spike timing information would continue to outperform rate-based decoders (Downer et al., 2021), since we commonly observed single cells whose responses supported sweep direction decoding well above the significance criterion in all cortical fields.

We compared decoding accuracy for the rate-based classifier for A1 against each of the nonprimary fields, for each sweep velocity (MATLAB kstest2). Direction selectivity did not differ significantly from A1 in any of the nonprimary fields we sampled (p > 0.01; uncorrected), likely due to floor effects given the universally poor performance of the rate-only classifiers. In summary, firing rate differences were rarely effective for discriminating sweep direction for all tested fields. The similarities among the distributions of decoding accuracy for the full spike train and timing-only classifiers demonstrate that spike timing carried nearly all the information about sweep direction in all tested cortical fields instead of overall firing rate.

To facilitate comparisons with prior studies that focused on rate-based analyses of FM sweep responses (i.e. Mendelson and Cynader, 1985?; Tian and Rauschecker, 2004), we also calculated a Direction Selectivity Index (DSI) that captured the difference in response strength to sweeps in different directions (see *Direction Selectivity Index (DSI)* in Methods). Overall, the prevalence of direction selectivity, as assessed by DSI values with absolute values greater than or equal to 0.33, indicating a relative doubling of responses in the preferred sweep direction was higher than the incidence of significant rate decoding, but still low: 60 ms: 9.5%; 120 ms: 12.1%; 180 ms: 11.5%; 240: 12.4%; 350 ms: 13.0%). Across all sweep velocities, DSI absolute values were less than 0.33 in 88.09% (4801/5450) of cases. If we consider DSI values with absolute values larger than 0.33, only 28.6% of them (182/637) corresponded to significant rate-based decoding, as determined by our methods. In 4.1% of cases (199/4801), significant rate decoding occurred for DSI magnitudes less than 0.33. These results suggest that DSI values do not unambiguously signal the presence of useful information about sweep direction since they do not account for trial-to-trial variability in cortical responses. Moreover, cortical neurons rarely exhibited consistent preferences for sweep direction across different sweep velocities. Results from a permutation test on the variance in DSI values across sweep speed indicated that neurons in our sample rarely (n = 9 out of 1090) exhibited lower variances that were significantly (p < 0.001) lower than chance (Only a single cell was significant for a criterion value of p < 0.05 Bonferroni corrected for 1090 tests).

As noted above, our goal in computing decoding accuracy for the full spike train, timing-only, and rate-only classifiers was to parse the contributions of spike timing and spike rate information to sweep discrimination in different auditory cortical fields. For example, a reduction in relative accuracy of the full spike train compared to the rate-only classifier in a belt field, relative to AI, would suggest a shift in how FM sweeps were encoded, and could evidence a transformation in how FM was represented. For each cortical field, we computed the population mean decoding accuracy, averaged over sweep velocity for each cell, obtained with the full spike train, timing-only, and rate-only classifiers for 3 sweep sets : 1) all sweeps (Figure 6); 2) sweeps in each cell’s better direction (Figure 7); 3) and matched-velocity pairs (i.e., direction selectivity; Figure 8). This produces a set of 9 values for each cortical field. Since chance accuracy varies across the different sweep sets, we normalized the values to the maximum obtained for the most informative cortical field and classifier to eliminate spurious correlations related to sweep counts.

In order to determine whether neurons in non-primary fields exhibit hierarchical transformations in the nature of FM sweep encoding, we computed the Pearson’s correlation coefficient between the patterns of decoding accuracy values between A1 and each non-primary field. If the encoding strategy remained similar, the correlation coefficient should be high, whereas if the coding strategy changed (e.g., from a temporal to a rate code) then the correlation coefficients should be low. The results of this analysis showed that the r-values exceeded 0.96 in all cases (CL: 0.97, p = 8.431e-06; CM: 0.96, p = 3.390e-05; ML: 0.99, p = 8.619e-08; MM: 0.98, p = 5.453e-06; R: 0.99, p = 2.819e-07), indicating that the form of FM sweep encoding appears to be highly conserved across all the tested fields, despite differences in decoding accuracy values. We performed an analogous analysis by computing the proportion of cells that exhibited decoding accuracy above chance for the same classifiers and sweep sets (without normalization, since the significance criteria were specific to each sweep set). Again, the A1 pattern was highly predictive of patterns in nonprimary fields (CL: 0.95, p = 7.692e-05; CM: 0.98, p = 6.220e-06; ML: 0.97, p = 1.194e-05; MM: 0.98, p = 1.370e-06; R: 0.95, p = 6.625e-05). In effect, while there are small but significant differences in how well nonprimary fields encode FM sweeps, there do not appear to be significant differences in how they encode them.

We also investigated the relationship between the absolute firing rates and the performance of the different classifiers to determine how firing rate constrains the informativeness of cortical spike trains. We counted all spikes within the analysis window used for spike train decoding (0-400 ms), then computed the (Pearson’s) correlation coefficient with decoding accuracy for each classifier. There was a significant positive correlation in A1 (r = 0.38; p = 1.017e-13), CL (0.40; 6.690e-08), CM (0.25; p = 3.595e-04), ML (0.47; p = 2.993e-10), and R (0.39; p = 1.216e-06). The only exception was MM (0.29; p = 0.36), where the correlation strength was similar but the significance was limited by the comparatively small sampled size.

In the foregoing analysis, the spikes we tallied were the spikes we decoded. We also quantified the responsiveness of each cell by counting the spikes within 200 ms of tone onset for a tone at BF across dB SPLs from 45 to 75 dB (in 10 dB increments). These stimuli were presented in a block of trials following the block containing the FM sweep stimuli randomly interleaved with tones at 65 dB SPL (see Methods). Even though none of these spikes contributed to the FM sweep responses that were decoded, the correlation values were similar to each other, and all were significant (A1: 0.49; p = 1.321e-23; CL: 0.37; p = 5.502e-07; CM: 0.39; p = 1.183e-08; ML: 0.47; p = 7.507e-10; MM: 0.36; p = 8.124e-03; R: 0.32; p = 9.415e-05). We repeated this analysis for decoding accuracies obtained with the timing-only classifier, and obtained highly similar values (A1: 0.49; p = 2.530e-23; CL: 0.35; p = 2.526e-06; CM: 0.43; p = 1.204e-10; ML: 0.48; p = 1.952e-10; MM: 0.40; p = 3.095e-03; R: 0.32; p = 2.701e-05), despite the firing rate normalization intrinsic to the process. This analysis shows that more responsive neurons are, on average, more informative when spike timing information is decoded. By contrast, these relationships were much weaker for the rate-only decoder (A1: 0.16; p = 1.907e-03; CL: 0.26; p = 7.047e-04; CM: 0.16; p = 0.022; ML: 0.20; p = 0.012; MM: 0.00; p = 0.989; R: 0.20; p = 0.0164). The lower correlation values likely reflect a restriction of range, since decoding accuracy values were poor for the rate-only classifiers.

We also tested whether there was a difference in the predictive validity of the absolute firing rate for decoding accuracy from A1 to the non-A1 fields. To do so, we performed a permutation test, using differences in the correlation values as the test metric. For all cortical fields, and for all classifiers, the actual difference in r-values was not larger than expected by chance for randomly permuted field labels (n = 10,000; p < 0.05), indicating that absolute firing rate was similarly predictive in the non-A1 fields. The results of the permutation tests were similar when the firing rates obtained from the FM sweep responses were used to obtain the correlations. These results support the idea that the form of FM sweep encoding is consistent in primary and non-primary fields, despite differences in its quality.

## Discussion

Our primary objective for this study was to address the lack of data on frequency modulation encoding by the core and belt fields of auditory cortex in alert macaque monkeys. Frequency modulation is critical for perception and discrimination of a variety of naturalistic stimuli, including speech and animal vocalizations, and much of that modulation can be captured by the parameters that define FM sweeps: spectral range, direction, and velocity. Auditory cortex appears to be necessary for the perception of time-varying stimuli, including FM sweeps, and the FM sweep spectral and temporal parameters used in this study encompass much of those parameters across encountered stimuli. Bilateral lesions of auditory cortex result in the inability for macaques to discriminate FM sweeps (Harrington et al., 2001), as well as between conspecific “coo” calls, which are characterized by FM components (Heffner and Heffner, 1984). Unfortunately, the ability to discriminate sweep velocity and direction (i.e., upward vs. downward) has been less well studied in intact animals. The ability to discriminate sweep direction was tested as a function of sweep speed in rats (Gaese et al., 2006) and cats (Zhang, 2010), with a similar result seen in gerbils using a different paradigm (Rybalka et al., 2006; Saldeitis et al., 2022). All of these studies found that discrimination performance was at or near peak performance for the sweep speeds used in the present study. Stimuli employed in studies in monkeys do not have perfect parallels with the stimuli used in these studies, but the FM speeds and directions we employed should be saliently different and discriminable by primate listeners (see Harrington et al., 2001). Thus, high decoding accuracy would be expected among a signification portion of neurons encountered in auditory cortex of the alert primate, as demonstrated here with full spike train and timing-only decoders.

In this study we had two main goals. The first was to better understand how single neuron responses in different cortical areas in alert primates encode FM sweep trajectories. The second was to compare FM sweep encoding by primary auditory cortex (A1) to that of non-primary, “downstream” cortical areas. Previous studies in these same monkeys (and, to a large extent, the same neurons) showed clear differences between A1 and area CL for spatially-varying stimuli (Woods et al., 2006) and that the responses across CL neurons could encode spatial location as a function of stimulus intensity consistent with psychophysical performance (Miller and Recanzone, 2009), making a strong case for a neural correlate of acoustic space perception. Here, by holding the spectral range constant we were able to investigate responses to stimuli that varied only in the time domain to ask whether a similar neural correlate could be revealed.

For the first goal, our strategy was to measure the information content of cortical responses to FM sweeps with nearest-neighbor Euclidean distance classifiers (Foffani and Moxon, 2004), as well as classifier variants that eliminated spike-timing information or normalized out spike rate information (Malone et al., 2007). Prior studies, particularly those conducted in anesthetized animals, typically focused on firing rate alone, and compared differences based on either overall rate or contrast sensitivity functions to determine single neuron selectivity for either direction or velocity (e.g. Mendelson and Cynader, 1985; Mendelson et al., 1993; Tian and Rauschecker, 2004). Consistent with those reports, we did find that our rate-based decoders, across the population, performed above chance (see Fig. 8), but the vast majority of neurons showed rate based decoding accuracy that was not statistically significant by our criteria.

Importantly, a neuron’s BF constrains the information it provides about FM sweep direction and velocity when spiking information is fully considered. This constraint can be ignored for rate coding models that ignore when spikes occur, provided that the FM sweeps span the hearing range. As we report here, however, firing rate information about FM sweep direction and velocity is quite impoverished in alert primate cortex in every cortical field we explored. Thus, while firing rates of primate auditory cortical neurons can provide sufficient information to discriminate space-varying stimuli, they do not appear to do so for the time-varying stimuli we investigated in this study in fields AI, CL, CM, ML, MM, or R. When spike timing was considered, decoding accuracy could be quite high in neurons distributed widely across auditory cortex. Moreover, removal of firing rate information by normalizing the PSTH typically had relatively little effect. Malone et al. (2017) had previously shown that in squirrel monkey auditory cortex, the removal of firing rate information slightly but significantly improved decoding of FM sweeps, indicating that the dominance of spike timing based encoding of FM sweeps appears to be consistent across primate species. These findings are consistent with a response model where the FM sweep interacts with the spectral tuning of a given cortical neuron to produce spikes at a particular latency with respect to sweep onset, rather than rate-based encoding of extracted FM features, such as direction, or velocity.

It is crucial to recognize that the decoder must have access to the stimulus onset time in order to exploit differences in response latency to discriminate among FM sweeps in different directions and velocities. The binned representation of spike trains fed to the classifiers embeds this latency information. The peak rate decoder (see Methods) represents the converse case—there, we identified a fixed window that maximized the spike count on a stimulus-by-stimulus basis, and used that window to feed the trial-by-trial spike count to the rate-only classifier. In our hands this procedure underperformed compared to a single, large fixed window for analysis (0-400 ms). This makes clear that even when rate-coding models are “tuned” to register spikes only during the window in which they occur, the omission of temporal information about the window itself (i.e., when it begins and ends), caused rate-decoding models to fail for this dataset.

As our results show, information about a neuron’s spectral tuning and sweep onset must be combined to interpret that neuron’s spiking output with respect to FM sweep trajectory. Thus, it is likely less useful to think of a given cortical neuron as encoding a particular direction or velocity in the abstract, rather than signaling a change in the acoustic energy in its frequency response area at some latency from the time of occurrence. Given a network-intrinsic reference signal (Panzeri et al., 2014), such as a frequency-independent onset response (Malone et al., 2015), it would be possible for downstream neurons to exploit spike timing information to explicitly represent FM sweep parameters. The decoding models we employed should not be understood as proxies for the readout mechanism that is causally upstream of perception. Instead, decoding accuracy quantifies a lower bound on the quality of FM sweep encoding (Schnupp et al., 2006). We used different classifier variants (full, rate-only, timing-only) to gauge the form of the encoding. Our data demonstrate that in awake primates, a temporal-to-rate conversion (i.e. Joris et al., 2004; Wang et al., 2005, 2008) has yet to occur for FM sweeps in the cortical fields we studied in this report.

Our second chief objective was evaluating evidence of hierarchical processing of FM sweeps. We expected clear differences in FM encoding between belt fields and A1, given previous results using vocalization stimuli (Rauscheker et al., 1995), human imaging studies (see Rauschecker and Afsahi, 2023; Martin et al., 2024) and spatial tuning (Woods et al., 2006; Miller and Recanzone, 2009). We focused on comparing A1 against five non-primary cortical areas for two reasons: The first was statistical robustness, since we had considerably more data from A1. The second was the fact that A1 is well connected with each of these five fields (Kaas and Hacket, 2000). We made every effort to reduce bias in these comparisons, including limiting the analysis to non-A1 neurons that had BFs matched to A1 neurons. There was no compelling evidence that non-A1 fields encode FM information qualitatively differently than A1 neurons do, nor was there any indication of the forms of hierarchical processing seen in the spatial domain (Recanzone et al., 2000b; Woods et al., 2006; Miller and Recanzone 2009; Engle and Recanzone 2013), where spatial tuning defined by the firing rate during stimulus presentation is sharpened between A1 and CL, providing enough spatial information to account for sound localization ability across stimulus intensities.

Our results largely contradict those of an earlier study of FM sweep processing in the lateral belt in the same species (macaque monkeys; Tian and Rauschecker, 2004). In the prior study, the authors reported that most neurons in the lateral belt were highly selective for the rate and direction of FM sweeps, and observed differences in the preferred FM rates between areas of the lateral belt. Notably, the DS metric calculated in Tian and Rauschecker (2004) was applied to each FM sweep velocity independently; a cell was considered ‘direction selective’ if any velocity crossed the criterion value (0.33). By contrast, we found that significant direction selectivity was rare. We reported the incidence of direction selectivity for each tested velocity, however, rather than considering a cell direction selective for a significant result at one of the 5 velocities we tested, which could explain some of the discrepancy. Further, DSI values exceeding the criterion value (0.33) did not exceed our decoding significance criterion in the majority (71%) of cases. Similarly, most cortical neurons did not exhibit rate-based preferences for sweep velocities, as evidenced by the fact that the median values for rate-based decoding accuracy were near chance performance in all cortical fields (Figure 6). Neither were there differences between belt fields and A1 that suggested increased rate-based selectivity for FM sweeps, nor tuning preferences specific to particular fields.

Given the contrast between this and the previous report, it is worth considering methodological differences between both studies in detail. First, the prior study used linear sweeps where the frequency range was dependent on the best frequency of the neuron understudy, as opposed to a constant frequency range as in the present study. Second, most previous studies, including Tian and Rauschecker (2004) involved the use of anesthetized rather than awake animals. This might explain why the results of that study were more consistent with prior studies of FM sweep processing in other anesthetized species, such as cats, by the same group (Tian and Rauschecker, 1994, 1998) and others (Mendelson and Cynader, 1985; Mendelson et al., 1993), where rate-based preferences for sweep direction and velocity were also commonly observed. Importantly, direction selectivity was assessed with the DSI (Mendelson and Cynader 1985; Heil et al. 1992a,b; Phillips et al. 1985; Shamma et al. 1993; Tian and Rauschecker 1994, 1998) but not accounting for the variance in the trial-by-trial responses or applying a statistical criterion. In the Tian and Rauschecker (2004) study the peak firing rate used to calculate the DSI was calculated for a 10 ms window slid across the averaged firing rate for that neuron at that stimulus speed and direction. The present study used a consistent 400 ms time window for all stimuli, justified by our finding that using the peak firing rate across tailored time windows did not improve performance over the 400 ms window. Further, Atencio et al. (2007), using this metric in awake owl monkeys, reported a median value of 0.04 across all neurons in A1, consistent with our results in the macaque.

As noted above, our decoding analyses indicated that the decoder was near chance in the vast majority of neurons when only using rate information. When we disaggregated FM sweep responses by sweep direction, (e.g., Figure 6), the decoding accuracy of the rate-only classifier indexes the quality of the representation of FM sweep velocity. The small number of points above the significance criterion in Figure 6 indicates the low prevalence of significant tuning to sweep velocity in our sample. Thus, there was little evidence of rate-based encoding of FM sweep features like direction or velocity, abstracted from sweep trajectory. We also observed very high (>0.96) correlations when we analyzed the patterns of relative decoding accuracy across all cortical fields, indicating that the transition from A1 to secondary fields was not accompanied by a shift from spike timing to rate-based representations of FM sweep features like direction and velocity.

In summary, non-primary fields were largely similar to A1 in encoding FM sweep trajectories via rapid changes in firing rates over the course of FM sweeps, rather than encoding abstracted FM sweep features as differences in average firing rates. The low incidence of direction selectivity and velocity tuning in our data reflects the fact that our classifier-based analyses uniformly applied a rigorous statistical significance criterion to qualify as selective or tuned. We think the most likely explanations for the higher incidence of tuning for FM features and differences across cortical fields are due to differential effects of anesthesia, the instability of rate estimates from narrow temporal windows (e.g., 10 ms), and differences in analytical methods, rather than field-specific processing differences.

The apparent lack of hierarchical specialization for processing FM sweep trajectories could be due to a number of factors. First, the FM features of sounds may not be as salient or ecologically relevant as spatial features, and their representations are not selected for transformation in the ascending auditory pathway. However, this seems unlikely this early in the processing pathway because stimulus features involving frequency modulation are ubiquitous in vocalizations and many other natural stimuli. An important difference between spectral and spatial acoustic features is the fact that in contrast to the visual system, it is the stimulus frequency, rather than the physical location, that is mapped to specific positions on the transduction surface for audition. As a result, central computation is required to infer the spatial positions of sounds, whereas spectral frequency information is immediately available. This difference could explain why the spectral features of sounds appear to persist in the responses of belt neurons (and neurons in field R), such that spike timing information related to FM sweep trajectory dominates their firing patterns to an extent very similar to primary auditory cortex. In other words, the fact that spectral frequency is fundamental to auditory processing might postpone the extraction of FM features along the auditory neuraxis. Thus, rate-based encoding of abstracted FM features may not emerge until higher order cortical areas such as the parabelt or more rostral areas of the temporal lobe. Further studies targeting those structures will be necessary to determine whether or not that is the case. The emergence of such hierarchical transformations is also likely to depend on ecological factors, such that earlier emergence (and explicit topographic mapping) might be expected in animals with highly specialized auditory systems, such as echolocating bats (see Suga, 2018), but not in auditory generalists like primates.

Below, we consider technical arguments that could explain the lack of belt specialization for the discrimination of FM sweep trajectories. First, one might argue that belt field neurons are disadvantaged by the inclusion criteria that we applied (see Methods), which are based on tone pips and FM sweeps, both of which are spectrally narrowband within a narrow time window. By this argument, the belt neurons included in this study would be more similar to A1 neurons than the broader population of belt neurons, since belt neurons that exhibited strong preferences for broad spectrum stimuli (Rauschecker et al., 1995; Rauschecker and Tian, 2004) would have been excluded. This argument lacks merit, since we included cells that responded in any significant way to either tones *or* the FM sweeps. Thus, if there were cells that responded *only* to spectrally broadband stimuli (i.e., noise or band-passed noise) but somehow contributed to the decoding and perception of FM sweeps, they would have been included here. Further, previous data in alert macaques show that many neurons in belt fields respond quite similarly to neurons in the core fields A1 and R (Recanzone et al., 2000a; Lakatos et al., 2005).

Next, we think it is unlikely that contralateral stimulus presentation disadvantaged belt neurons by excluding neurons selective for a different location than directly opposite the contralateral ear, since even the sharply tuned neurons in CL have some significant response across virtually all locations in azimuth, and certainly all in contralateral space, with the peak of the population response exactly where the stimuli of this study were presented (see Woods et al., 2006). Further, neurons in ML and R showed equivalent spatial tuning to those of A1, which is where we predicted differences would be most apparent. Thus, we think our population of non-A1 neurons is broadly representative of non-A1 neurons in this preparation.

One might also argue that belt neurons could have been disadvantaged by the use of Euclidean distance nearest neighbor classifiers, as opposed to more sophisticated techniques. We cannot definitively rule out the possibility that there is a readout mechanism that could extract more information from belt neurons than from A1 neurons, resulting in higher decoding accuracies and supporting claims about specialization for extracting FM sweep parameters in higher-order cortex. Nevertheless, we believe that using simple, interpretable classifiers based on the full spike train, rate-only, and timing-only features of cortical spike trains on individual trials provides a good estimate of the information available to higher order cortical areas, and is appropriately unbiased. It is also possible that evidence for hierarchical specializations for FM sweep processing is available in the population responses of belt neurons, which may require specific readout mechanisms to extract all the available information via rate codes (e.g., Johnson et al., 2025).

Finally, we consider whether hemispheric differences could explain the lack of specialization for FM sweep processing we observed. Our recordings were exclusively performed in the left hemisphere of the three monkeys, making that consistent with data on FM selectivity in primates from other groups (rhesus monkey: Tian and Rauschecker, 2004; owl monkey: Atencio et al., 2007; squirrel monkey: Godey et al., 2005; marmoset: Kajikawa et al., 2008) as well as those from other species (ferret: Shamma et al., 2993; cat: Phillips et al., 1985; Heil et al, 1992a,b). This allows us to make direct comparisons between species. It should be noted that lesion studies in gerbils indicate that lesions of the right hemisphere produce larger deficits in FM direction discrimination (Rybalka et al., 2006; Wetzel et al., 2008; see also Saldeitis et al., 2022). This would suggest that we may have seen either greater discrimination across fields, or hierarchically between fields if we had studied the right hemisphere instead of the left. However, a study in squirrel monkey recorded from both hemispheres and found no differences (Godey et al, 2005). Additionally, patients with left hemisphere PT lesions did show significant deficits in FM sweep direction discrimination (Martin et al., 2024). Thus, it remains to be seen if there are hemisphere differences for these parameters in macaques.

In summary, we observed remarkable consistency in the nature of the representation of FM sweep trajectories from primary to non-primary areas in the rhesus auditory cortex. In particular, we observed significant variance in decoding accuracy in all tested fields, and, notably, failed to observe a ‘ceiling’ on decoding accuracy in nonprimary fields, despite a small but significant reductions in median accuracy (excluding MM) relative to A1, which boasts superior temporal resolution relative to R (Scott et al., 2009) and most belt fields (Recanzone 2000a). Our results suggest that further studies of tertiary cortical fields such as the parabelt (i.e. Kajikawa et al., 2015) will be necessary to understand the contribution of the rostral core and belt fields to higher-order auditory processing in the temporal domain. Finally, while auditory spatial processing has shown clear similarities to the ‘where’ processing stream of visual cortex, the ‘what’ or ‘non-spatial’ processing analog has remained more elusive. This may well be that temporal processing is ubiquitous across all levels of auditory processing given that the stimuli that need to be processed are inherently continually time-varying.

## Acknowledgments

The authors would like to thank Timothy Woods, Steve Lopez, James Long, Joanne Rahman, Daniel Seto and Nathan Beckerman for their participation in the collection of the single neuron data, and the California National Primate Research Center for expert animal care.

## Arthur Contributions

GHR conceived and designed the original research and data collection.

BJM analyzed the data with input from GHR.

GHR performed the experiments and oversaw others that collected the data (see ‘acknowledgements).

BJM and GHR interpreted the results of the experiments / analysis.

BJM prepared the original figures which were revised by BJM and GHR

BJM and GHR jointly drafted, edited and revised the manuscript.

BJM and GHR approved the final version of the manuscript.

